# Ancestral SARS-CoV-2 immune imprinting persists on RBD but not NTD after sequential Omicron infections

**DOI:** 10.1101/2024.05.30.596664

**Authors:** Zuowei Wang, Ling Li, Ruiping Du, Xixian Chen, Yi Sun, Rongrong Qin, Yunjian Li, Hualong Feng, Lin Hu, Xuanyi Chen, Maosheng Lu, Liwei Jiang, Teng Zuo

## Abstract

Whether Omicron exposures could overcome ancestral SARS-CoV-2 immune imprinting remains controversial. Here we analyzed B cell responses evoked by sequential Omicron infections in vaccinated and unvaccinated individuals. Plasma neutralizing antibody titers against ancestral SARS-CoV-2 and variants indicate that immune imprinting is not consistently induced by inactivated or recombinant protein vaccines. However, once induced, immune imprinting is not countered by successive Omicron challenges. We compared binding specificities, neutralizing capacities, developing origins and targeting epitopes of monoclonal antibodies from individuals with or without immune imprinting. Although receptor-binding domain (RBD) and N-terminal domain (NTD) of spike are both primary targets for neutralizing antibodies, immune imprinting only shapes antibody responses to RBD by impeding the production of Omicron-specific neutralizing antibodies while facilitating the development of broadly neutralizing antibodies. We propose that immune imprinting can be either neglected by NTD-based vaccines to induce variant-specific antibodies or leveraged by RBD-based vaccines to induce broadly neutralizing antibodies.

## Introduction

Immunological memory is a hallmark of adaptive immunity, which consists of humoral and cellular immunity(1). Upon primary exposure to an antigen, humoral immune responses typically start from priming of naive B cells through cognate interaction between B cell receptor (BCR) and antigen in follicles of secondary lymphoid organs(2). Primed B cells migrate to the border between B cell follicle and T cell zone, where they encounter activated CD4 T cells. Cognate T-B interaction leads to clonal expansion of B cells and formation of germinal centers (GCs). Within GCs, B cells undergo somatic hypermutation (SHM) on variable region of BCR to generate a pool of mutated B cells, which are then selected based on their affinities to the antigen. After repeated rounds of mutation and selection, affinity-matured GC B cells differentiate into plasmablasts (PBs), plasma cells (PCs) or memory B cells. PB/PCs constantly secrete antibodies and maintain high levels of serum antibodies for months to years. Meanwhile, memory B cells circulate throughout the body until they encounter cognate antigens. Activated memory B cells rapidly differentiate into PB/PCs to secrete antibodies, or to a lesser extent, into GC B cells to undergo another round of affinity maturation(3).

As the basic principle of long-term immune protection against pathogens after infection or vaccination, immunological memory ensures much faster and stronger immune responses upon antigen re-exposure(1). However, the effects of immunological memory could be undesirable when antigens in prior and subsequent exposures are heterologous but antigenically related. One example of such scenario is that individuals’ antibody responses against influenza viruses are largely biased towards the predominant strains in their early lives(4). This phenomenon is referred to as original antigenic sin or immune imprinting, which describes the effects of immunity elicited by prior antigen exposures on subsequent immune responses to antigenically related antigens.

Since the outbreak of COVID-19 pandemic, the ancestral SARS-CoV-2 has kept evolving and generated hundreds of thousands of variants. Among them, Omicron emerged in November 2021 and accumulated more than 30 mutations on spike, the target of neutralizing antibodies(5). Over the past two and a half years, the predominant strains have been rapidly iterated by Omicron-derived variants, including BA.1, BA.2, BA.4, BA.5, BQ.1, XBB.1.5, XBB.1.16, XBB.1.9, EG.5.1 and BA.2.86.

These variants exhibit unprecedented neutralization evasion from antibodies elicited by ancestral SARS-CoV-2 infection or vaccines based on ancestral SARS-CoV-2 (6–11). Consequently, several waves of infection by Omicron variants have occurred and updated vaccines have been developed to match those variants (12–14).

Analysis of B cell responses in individuals who experienced breakthrough infection of Omicron variants or received Omicron booster vaccines demonstrates the induction of immune imprinting by ancestral SARS-CoV-2(15–20). Moreover, the imprinted antibody responses display substantially reduced neutralizing activities against later Omicron variants compared with ancestral SARS-CoV-2 and earlier Omicron variants, highlighting that immune imprinting may impede the generation of neutralizing antibodies against future SARS-CoV-2 variants or other related viruses(21). Notably, a recent study reported that sequential Omicron exposures could overcome immune imprinting and induce a large portion of *de novo* Omicron-specific antibodies(22). In contrast, other studies concluded that a single exposure to antigenically distant XBB.1.5 or double exposures to Omicron variants rarely elicited Omicron-specific antibody responses in the presence of immune imprinting(23–25). Therefore, it is controversial whether Omicron exposures could override ancestral SARS-CoV-2 immune imprinting.

In this study, we compared human B cell responses elicited by sequential Omicron infections in the presence or absence of ancestral SARS-CoV-2 immune imprinting. By measuring plasma neutralizing antibody titers against ancestral SARS-CoV-2 and variants, we find that immune imprinting is not consistently induced by inactivated or recombinant protein vaccines. However, once induced, immune imprinting is not countered by repeated Omicron exposures. Comprehensive analysis of monoclonal antibodies from individuals with or without immune imprinting demonstrates that RBD and NTD are both primary targets for neutralizing antibodies, whereas only RBD-specific antibody responses are imprinted. Consequently, Omicron-specific neutralizing antibodies to RBD are hindered while broadly neutralizing antibodies to RBD are elicited. These findings provide new insights into the development of next-generation SARS-CoV-2 vaccines.

## Results

### Sequential Omicron infections could not overcome ancestral SARS-CoV-2 immune imprinting induced by inactivated or recombinant RBD vaccines

To examine whether Omicron exposures could overcome ancestral SARS-CoV-2 immune imprinting, we collected peripheral blood from 12 donors of sequential Omicron infections (Figure 1A). They were first infected in December 2022 or January 2023, when BA.5 and BF.7 were predominant strains(26). Around half a year later, they were infected again when XBB lineages (XBB*) were predominant strains(27). Before the infections, Donor A and B were not administrated with any vaccines based on ancestral SARS-CoV-2, while the other donors received two or three doses of inactivated vaccine or recombinant RBD vaccine or in combination.

**Figure 1.**
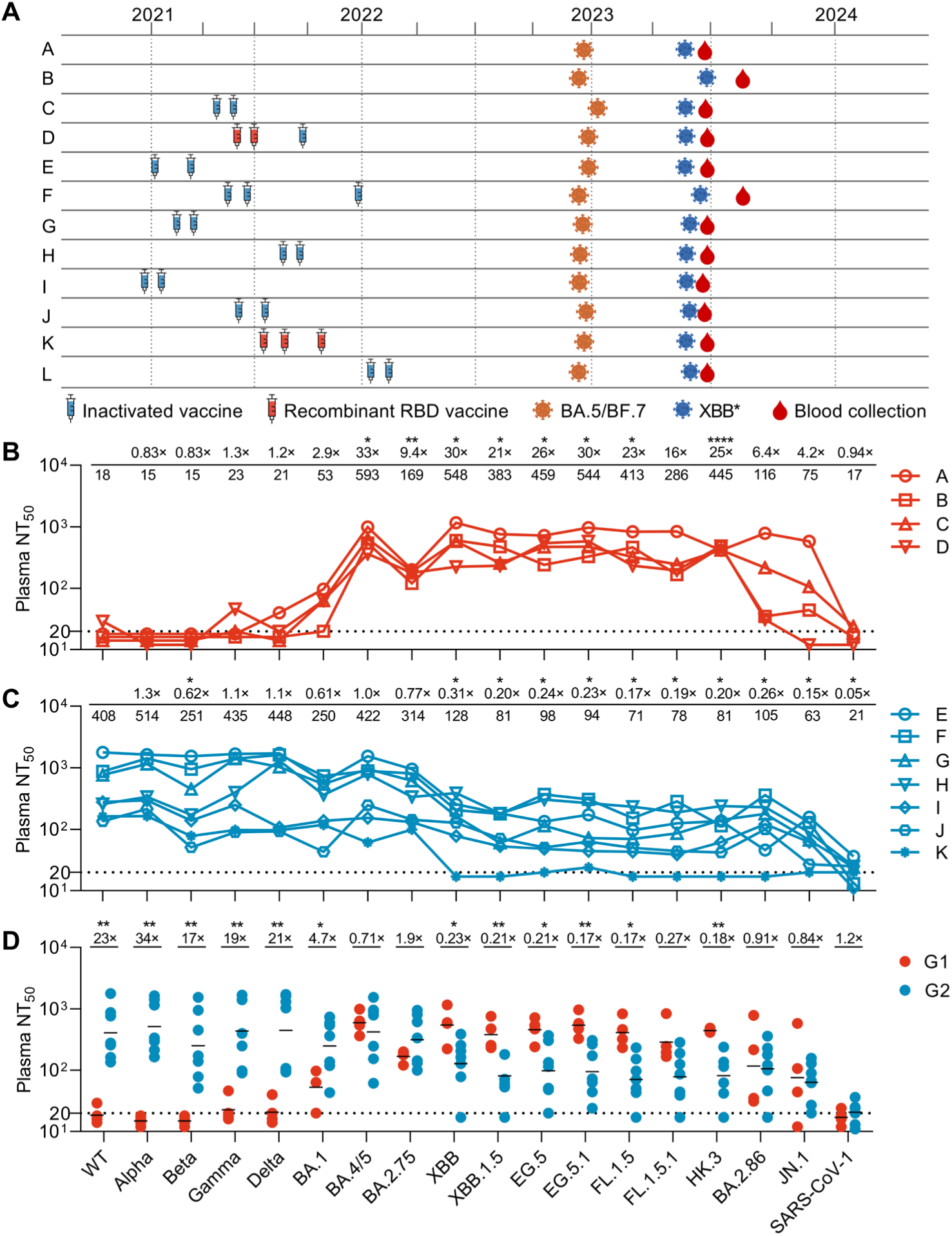
Sequential Omicron infections could not overcome ancestral SARS-CoV-2 immune imprinting induced by inactivated or recombinant RBD vaccines. (A) Vaccination and infection information of 12 donors. (B, C) Summary of plasma neutralizing antibody titers (NT50). For plasma with no neutralizing activity at 20-fold dilution, a number between 10–20 is given as titer to separate the curves. The numbers below the line are geometric mean titers against the pseudoviruses. The titers against WT are compared with other variants and the fold changes are indicated above the line. For panel B, statistical analysis was performed by two-tailed paired T test. For panel C, statistical analysis was performed by two-tailed Wilcoxon matched-pairs signed rank test. *P < 0.05, **P < 0.01, **** P < 0.0001. (D) Comparison of neutralizing antibody titers of G1 (Donor A-D, unimprinted group) and G2 (Donor E-K, imprinted group). The numbers on top are fold changes. Statistical analysis was performed by two-tailed unpaired T test. *P < 0.05, **P < 0.01. The neutralizing titers are calculated with results from two independent experiments, in which duplicates are performed.

We measured plasma neutralizing antibody titers against a panel of 18 pseudoviruses bearing spikes from ancestral SARS-CoV-2 (WT), Alpha, Beta, Gamma, Delta, BA.1, BA.4/5, BA.2.75, XBB, XBB.1.5, EG.5, EG.5.1, FL.1.5, FL.1.5.1, HK.3, BA.2.86, JN.1 and SARS-CoV-1 (Figure S1A). Donor A-D show poor or lower neutralizing titers against earlier SARS-CoV-2 strains from WT to BA.1 while higher titers against strains from BA.4/5 to HK.3 (Figure 1B). As Donor A and B were unvaccinated, their patterns represent antibody responses induced by sequential Omicron exposures in the absence of immune imprinting. Despite the fact that Donor C and D were vaccinated, they display similar patterns with Donor A and B, suggesting that immune imprinting is not induced by vaccination in Donor C and D. Donor E-K maintain higher titers against earlier strains from WT to BA.2.75 while lower titers against XBB and later strains, indicating the presence of immune imprinting in these individuals (Figure 1C). Different from above donors, Donor L exhibits a quite chaotic pattern of neutralizing titers and thus is excluded for further analysis (Figure S1B).

We compared neutralizing titers of Donor A-D (referred to as unimprinted group, G1) and Donor E-K (referred to as imprinted group, G2) (Figure 1D). For early strains from WT to BA.1, imprinted group has significantly higher titers than unimprinted group. On the contrary, for more recent Omicron variants, including XBB, XBB.1.5, EG.5, EG.5.1, FL.1.5 and HK.3, unimprinted group surpasses the imprinted group. Taken together, these data suggest that immune imprinting is not consistently induced by vaccination. However, once induced, immune imprinting is not countered by sequential Omicron infections.

### Immune imprinting leads to more frequent cross-reactive antibodies to RBD but not NTD

To decipher the effects of immune imprinting on human B cell responses elicited by sequential Omicron infections, we moved on to isolate and characterize monoclonal antibodies (mAbs) from those donors. As they were sequentially exposed to BA.5/BF.7 and XBB*, we sorted single memory B cells or plasmablasts with XBB.1.5 spike as bait (Figure S2). After cloning antibody variable regions from single cells into corresponding antibody-expressing vectors, we transfected HEK293T cells with paired plasmids and collected supernatant for ELISA and neutralization assay. The supernatant was tested against spike (S), S1, RBD, NTD and S2 of XBB.1.5 and WT by ELISA, and tested against XBB.1.5 and WT pseudoviruses by neutralization assay (Figure 2A). In total, we identified 364 clones positive for XBB.1.5 spike from Donor A, B, C, D, E, F, G, I, J and K. The proportions of antibodies against each domain vary in a wide range and no significant differences are observed between unimprinted group (G1) and imprinted group (G2) (Figure 2B). With 50% as cutoff for neutralization of supernatant, 84 clones were identified as neutralizing antibodies against XBB.1.5 or WT or both (Figure 2A, B). Notably, cross-neutralizing antibodies are all from imprinted group (Figure 2A). In addition, the vast majority of neutralizing antibodies target either RBD or NTD and there are no significant differences between the two groups (Figure 2B).

**Figure 2.**
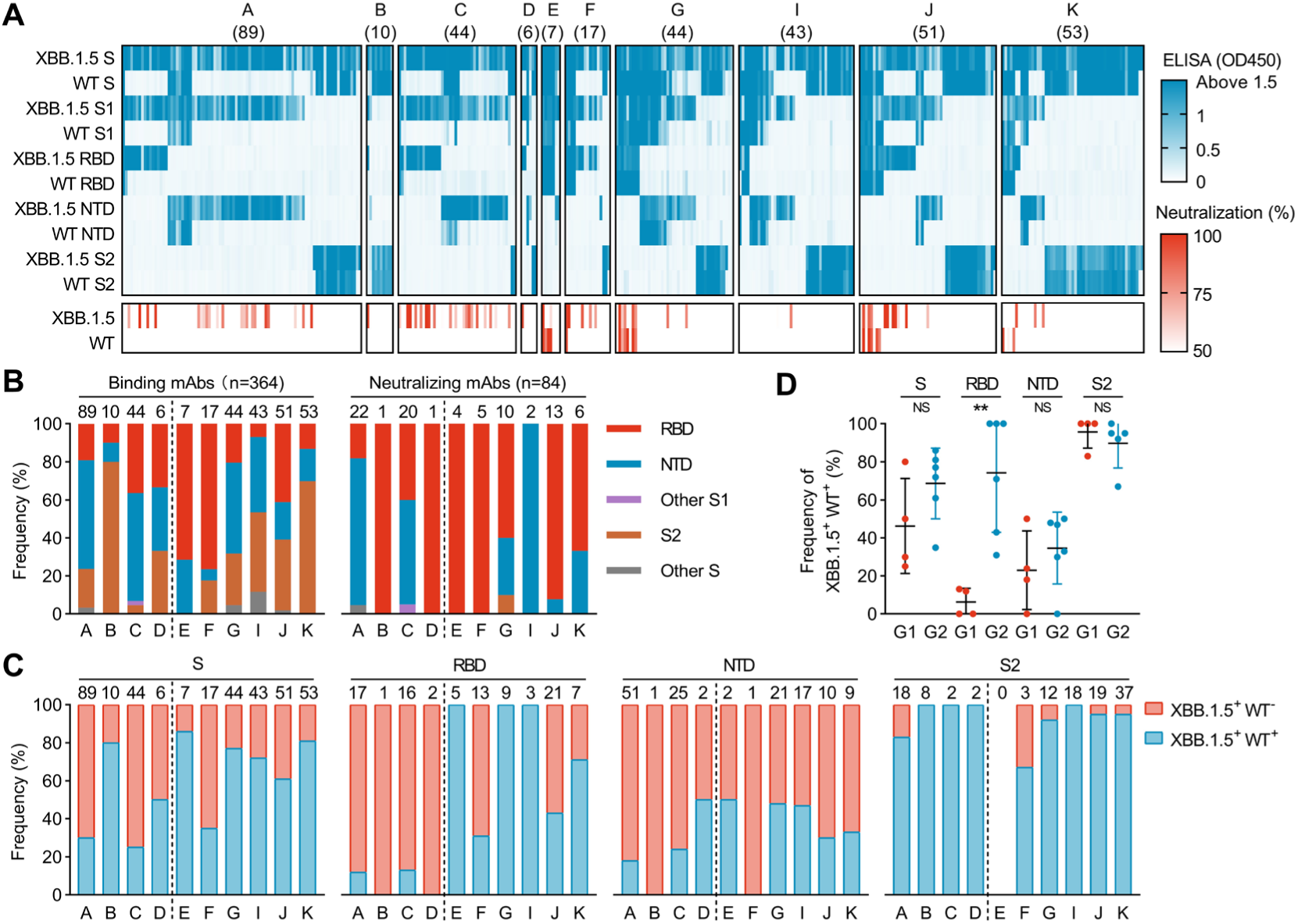
Immune imprinting leads to more frequent cross-reactive antibodies to RBD but not NTD. (A) ELISA and neutralization results of 364 mAbs. Supernatant of HEK293T cells was tested against Spike, S1, S2, NTD and RBD of WT and XBB.1.5 by ELISA, and tested against WT and XBB.1.5. pseudoviruses by neutralization assay. For ELISA, supernatant was not diluted. For neutralization, supernatant was diluted by 4.5-fold. The results are mean values of two independent experiments. For neutralization, duplicates are performed in each experiment. (B) Frequency of binding mAbs and neutralizing mAbs against different domains in each individual. (C) Frequencies of WT-cross-reactive antibodies (XBB.1.5^+^ WT^+^) and Omicron-specific antibodies (XBB.1.5^+^ WT^-^) in each individual. (D) Comparison of WT-cross-reactive antibodies between G1 and G2. Data are represented as mean ± SD. Statistical analysis was performed by two-tailed unpaired T test. NS (Not significant) p > 0.05, **P < 0.01.

We further analyzed the frequency of cross-reactive antibodies (XBB.1.5^+^ WT^+^) (Figure 2C, D). Most S2-specific antibodies are cross-reactive, which should be due to the conservation of this domain. For NTD-specific antibodies, the frequencies range from 0 to 50% and there is no difference between G1 and G2. In contrast, for RBD-specific antibodies, G2 has remarkably higher frequency than G1, which represents recall of cross-reactive memory B cells induced by vaccination. Overall, these results indicate that immune imprinting mainly influences antibody responses to RBD after sequential Omicron infections.

### Immune imprinting facilitates the development of broadly neutralizing antibodies against RBD

To determine the effects of immune imprinting on neutralizing antibodies, we purified RBD- or NTD-specific mAbs with supernatant neutralization above or close to 50%. A total number of 87 mAbs, including 16 RBD-specific and 32 NTD-specific mAbs from G1 and 33 RBD-specific and 6 NTD-specific mAbs from G2, were tested against a panel of SARS-CoV-2 pseudoviruses (Figure 3A). Similar to the neutralization patterns of plasma from G1, RBD-specific mAbs from G1 (G1-RBD) poorly neutralize early strains from WT to BA.1 while potently neutralize later Omicron variants. RBD-specific mAbs from G2 can be divided into two subgroups according to their neutralization patterns. One subgroup resembles mAbs from G1-RBD in neutralization pattern and is named as non-WT-neutralizing (Abbreviated as non-WT-neut). The other subgroup replicates the neutralization patterns of plasma from G2 and is termed as WT-neutralizing (Abbreviated as WT-neut). It is worth mentioning that there are several broadly neutralizing antibodies in this group, including KXD-E1, KXD-F3, KXD-F1, KXD-G10 and KXD-G5. Most NTD-specific mAbs are able to neutralize BA.4/5, BQ.1, XBB, XBB.1.5, EG.5.1 and HK.3, and the patterns are similar between G1 and G2. Regarding to the neutralizing potency (IC50s) against different strains, mAbs from G1-RBD are more potent to BA.4/5 than later variants whereas mAbs from G2-RBD-non-WT-neut exhibit similar neutralizing activity against BA.4/5 and later strains (Figure S3A). Antibodies from G2-RBD-WT-neut have equivalent potency to early strains from WT to BA.4/5 while decreased potency to later variants (Figure S3A). Different from RBD-specific mAbs, NTD-specific mAbs from both G1 and G2 are most potent to XBB and XBB.1.5 (Figure S3B).

**Figure 3.**
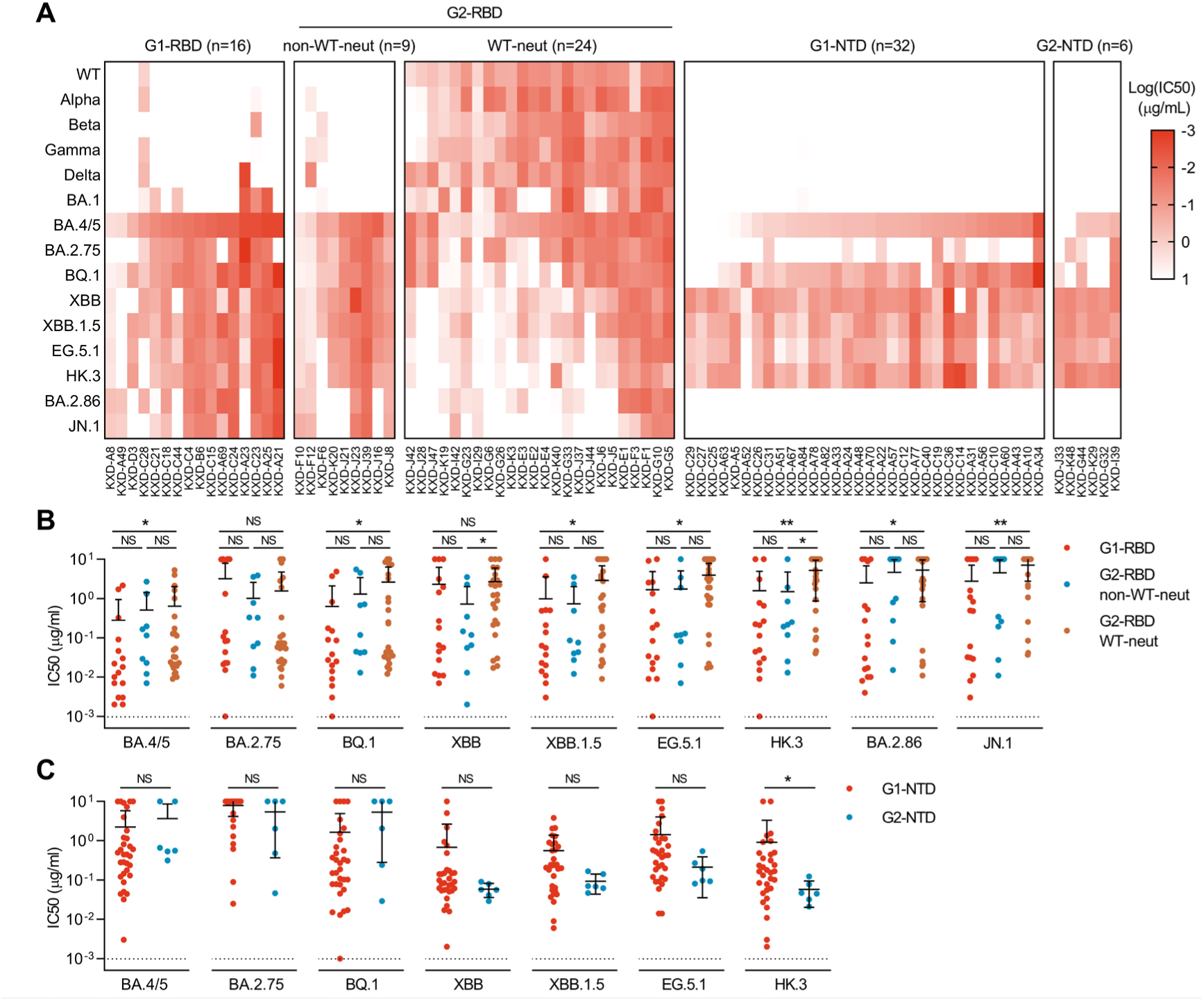
Immune imprinting facilitates the development of broadly neutralizing antibodies against RBD. (A) IC50s of mAbs against WT, Alpha, Beta, Gamma, Delta, BA.1, BA.4/5, BA.2.75, BQ.1, XBB, XBB.1.5, EG.5.1, HK.3, BA.2.86 and JN.1. The highest antibody concentration used to determine IC50 is 10μg/ml. (B) Comparing IC50s of RBD-specific mAbs from three groups. (C) Comparing IC50s of NTD-specific mAbs from two groups. Data are represented as the mean ± SD. Statistical analysis was performed by two-tailed unpaired T test. NS (Not significant) p > 0.05, *P < 0.05, **P < 0.01. IC50s are calculated with results from two independent experiments, in which duplicates are performed.

We compared the neutralizing potency of mAbs from different groups against the same strain (Figure 3B,3C). Overall, mAbs from G1-RBD have similar neutralizing potency with mAbs from G2-RBD-non-WT-neut. However, mAbs from G1-RBD are more potent than mAbs from G2-RBD-WT-neut against variants from BA.4/5 to JN.1 except BA.2.75 and XBB. On the other hand, NTD-specific mAbs from both groups have analogous potency against all tested strains except HK.3. These data further confirm that antibody responses to RBD while not NTD are affected by immune imprinting. Moreover, the imprinted RBD-specific antibodies tend to have broadly neutralizing activity although they generally show suboptimal neutralizing potency against Omicron variants.

### Repeated Omicron infections induce de novo Omicron-specific antibody responses in the presence of immune imprinting

To examine whether repeated Omicron infections could induce de novo antibody responses from naïve B cells in the presence of immune imprinting, we move on to investigate the origins of neutralizing mAbs from each group. In published literatures, it is generally assumed that cross-reactive antibodies derive from memory B cells induced by WT whereas omicron-specific antibodies originate from naïve B cells activated by Omicron(22–25). However, these assumptions remain to be confirmed by experiments to exclude possibilities that memory B cells induced by WT may lose cross-reactivity and Omicron-specific naïve B cells may develop cross-reactivity during affinity maturation. Here, we constructed germline versions of representative mAbs from each group and tested their binding to a panel of spikes expressed on HEK293T cells by FACS (Figure 4A). Germline antibodies from G1-RBD, G2-RBD-non-WT-neut, G1-NTD and G2-NTD bind Omicron spikes but not WT spike. Oppositely, germline antibodies from G2-RBD-WT-neut mostly recognize WT spike but not Omicron spikes. These data confirm that mAbs from G1-RBD, G2-RBD-non-WT-neut, G1-NTD and G2-NTD are de novo antibodies induced by Omicron, whereas mAbs from G2-RBD-WT-neut are recalled antibodies initially induced by WT.

**Figure 4.**
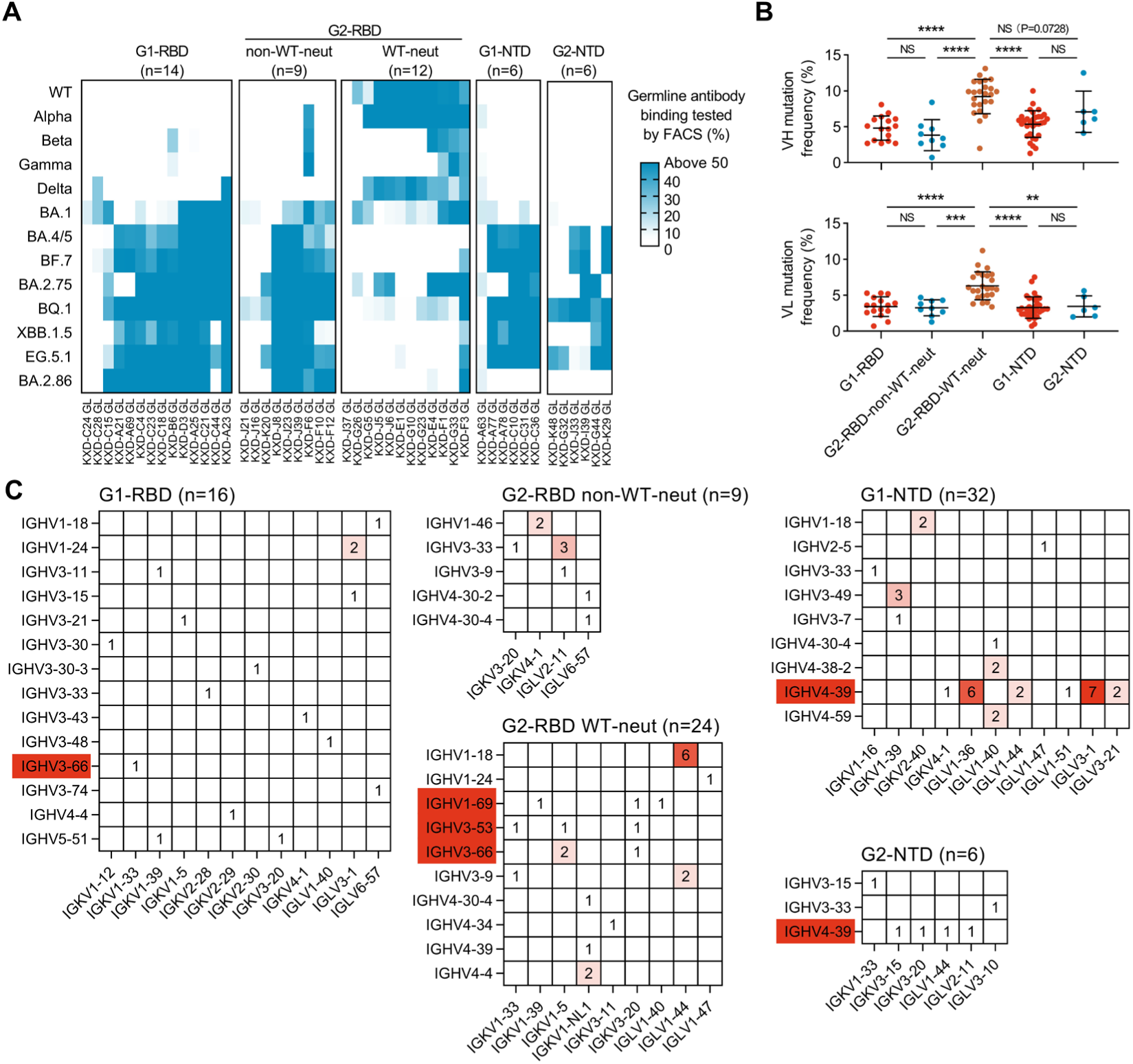
Repeated Omicron infections induce de novo Omicron-specific antibody responses in the presence of immune imprinting. (A) Binding specificities of germline antibodies measured by FACS. HEK293T cells expressing SARS-CoV-2 spikes were incubated with 50μg/ml germline antibodies and analyzed by FACS. The percentages of positive cells are calculated. The results are mean values of two independent experiments. (B) Somatic hypermutations of VH and VL. DNA sequences were analyzed. Data are represented as mean ± SD. Statistical analysis was performed by two-tailed unpaired T test. NS (Not significant) p > 0.05, **P < 0.01, ***P < 0.001, ****P < 0.0001. (C) Frequencies of V genes in each group. Frequently used V genes are highlighted by colored background.

Correlated with their origins, mAbs from G2-RBD-WT-neut show higher mutation levels than other mAbs, which reflects extensive affinity maturation sequentially driven by WT and Omicron (Figure 4B). On the other hand, we also analyzed the repertoire of each group (Figure 4C and Table 1). In G2-RBD-WT-neut group, IGHV1-18 and IGLV1-44 are most frequent as a consequence of B cell clonal expansion in Donor J. Moreover, IGHV3-53, IGHV3-66 and IGHV1-69, which have been identified as frequent VH genes for RBD-specific antibodies from individuals exposed to WT spike(28, 29), are frequently used in G2-RBD-WT-neut but not in G1-RBD and G2-RBD-non-WT-neut. For NTD-specific antibodies, IGHV4-39 is most frequently used and is identified from multiple donors in both groups. Therefore, these IGHV4-39 antibodies can be taken as a class of public antibody against NTD. In sum, these data suggest that repeated Omicron exposures could induce de novo antibody responses to RBD and NTD in the presence of immune imprinting.

### Omicron-specific and WT-cross-reactive antibodies recognize similar epitopes

To determine whether de novo antibody responses induced by repeated Omicron infections recognize distinct epitopes compared with WT-cross-reactive antibodies, we performed competition ELISA (Figure 5A, 5B). Three broadly neutralizing antibodies targeting RBD, including BD55-1205, BD56-1854 and BD55-5483 were taken as reference(30). A major cluster is formed by the majority of RBD-specific antibodies. Most antibodies in this cluster also compete with BD55-1205, BD56-1854 and ACE2. Outside the major cluster, a minor cluster is formed by 6 antibodies. They compete with BD55-5483 but not with BD55-1205, BD56-1854 and ACE2. The rest 3 RBD-specific antibodies do not exhibit apparent competition with other antibodies or ACE2. On the other hand, NTD-specific antibodies show competition with each other and form a single cluster, suggesting that NTD-specific neutralizing antibodies target a supersite. Taken together, these data provide explanation for the suppressed Omicron-specific antibody responses to RBD in the presence of immune imprinting.

**Figure 5.**
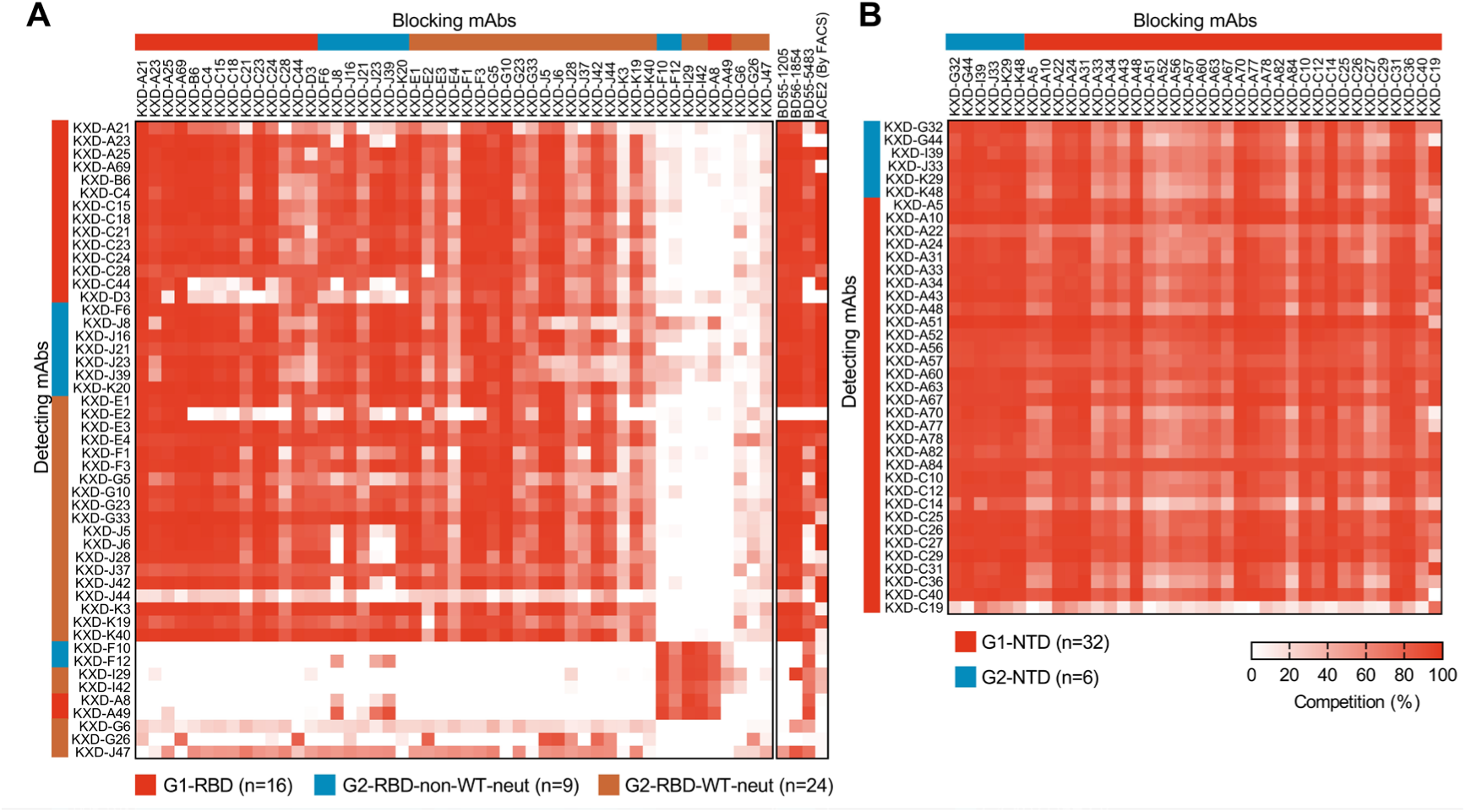
Omicron-specific and WT-cross-reactive antibodies recognize similar epitopes. (A) Competition ELISA of RBD-specific mAbs. Competition with ACE2 was performed by FACS. (B) Competition ELISA of NTD-specific mAbs. The results are mean values of two independent experiments.

## Discussion

In this study, we compared B cell responses evoked by sequential Omicron infections in individuals vaccinated or unvaccinated with inactivated or recombinant RBD vaccines. Unexpectedly, two vaccinees show no hints of immune imprinting, according to their polyclonal and monoclonal antibody responses against ancestral SARS-CoV-2 and variants. Therefore, they are classified into unimprinted group along with two unvaccinated individuals throughout this study. Considering the factor of individual variations, we believe that this classification is more reasonable than simply grouping the cohorts based on antigen exposure history, especially for cohorts with limited size. We speculate that no immune imprinting in these two individuals is probably due to weak immune responses stimulated by vaccination. Moreover, whether this phenomenon is common remains to be determined with larger cohorts.

In contrast to individuals without immune imprinting, individuals with immune imprinting have lower plasma neutralizing antibody titers against multiple Omicron variants after XBB, along with more frequent WT-cross-reactive antibodies and less frequent Omicron-specific antibodies to RBD. Therefore, we conclude that immune imprinting induced by inactivated or recombinant RBD vaccines is not countered by sequential Omicron infections, which is inconsistent to a study performed by Yisimayi et al(22). We suppose this contradiction is due to following factors. First, the infections were sequentially caused by BA.1/BA.2 and BA.5/BF.7 in their study whereas by BA.5/BF.7 and XBB* in our study. We notice that individuals with or without immune imprinting have equivalent plasma neutralizing antibody titers to BA.4/5 and BA.2.75. Moreover, individuals with immune imprinting show unaffected neutralizing titers against early Omicron variants including BA.1, BA.2.75 and BA.4/5 compared with WT. These data suggest that immune imprinting does not compromise antibody responses to early Omicron variants after repeated Omicron exposures. Second, all vaccinees were considered as imprinted in their study while two vaccinees were taken as unimprinted in our study. As vaccinees without immune imprinting are able to generate abundant Omicron-specific antibodies, excluding them from imprinted cohort may lead to more clear results.

A major finding of our study is that antibody responses to NTD are not affected by immune imprinting. As immunological memory and cross-reactivity are the basis of immune imprinting(31), one possible explanation for this finding is that NTD is less immunogenic than RBD because of its extensive glycan shielding(32, 33), and thus few NTD-specific memory B cells are induced. Another possible explanation is that NTD-specific memory B cells induced by WT are not cross-reactive to Omicron variants. However, consistent with previous studies(34–39), our results argue that NTD is highly immunogenic and able to elicit antibodies cross-reactive to XBB.1.5 and WT, which meet the preconditions for immune imprinting. Therefore, the mechanisms for the unimprinted antibody responses to NTD remain to be explored.

Similar to previous studies(15–18, 20, 22–24), we prove that immune imprinting persistently shapes antibody responses to RBD. As a result, RBD-specific antibodies can be divided into two groups: WT-cross-reactive antibodies and Omicron-specific antibodies. Compared with Omicron-specific antibodies, WT-cross-reactive antibodies tend to have broader neutralization spectrum. For example, KXD-F1 neutralize all tested pseudoviruses from WT to JN.1 with IC50s ranging from 0.006 to 0.084 μg/ml. As germlines of those WT-cross-reactive antibodies generally have limited breadth, there is no doubt that the breadth of WT-cross-reactive antibodies is attributed to affinity maturation. However, the underlying mechanisms are not fully understood. According to the model of affinity maturation in GCs, it is possible that WT-cross-reactive memory B cells become Omicron-specific after affinity maturation. By comparing the binding specificities of paired germline and mature antibodies, we find that all WT-reactive germline antibodies consistently recognize WT spike after affinity maturation, which negate the possibility like our previous study(40). Oppositely, it is also possible that Omicron-specific naïve B cells may become cross-reactive after affinity maturation. However, we have not found evidence to support the possibility, despite that fact that infection or vaccination with WT could elicit broadly neutralizing antibodies against Omicron variants(30, 41–43). More studies are required to solve these puzzles on affinity maturation.

Currently, ancestral SARS-CoV-2 immune imprinting has been acknowledged as a great barrier for vaccine update. According to our findings, we propose that NTD-based vaccines could be designed to bypass immune imprinting, since NTD is also a major target of neutralizing antibodies. On the other hand, vaccines containing RBD could be designed to leverage the potential of immune imprinting in eliciting broadly neutralizing antibodies.

## Materials and methods

### Ethics statement

This study was approved by the Ethics Committee of Hefei Institutes of Physical Science, Chinese Academy of Sciences (Approval Number: YXLL-2023-47). All donors provided written informed consent for collection of information, analysis of plasma and PBMCs, and publication of data generated from their samples.

### Human samples

Peripheral blood samples were collected from 12 donors (Figure1A). Plasma and peripheral blood mononuclear cells (PBMCs) were separated from blood by Ficoll density gradient centrifugation.

### Cell lines

HEK293T cells expressing human ACE2 (HEK293T-hACE2) were kindly provided by Prof. Ji Wang at Sun Yat-Sen University. HEK293T cells were from ATCC. HEK293T and HEK293T-hACE2 cells were cultured in DMEM with 10% fetal bovine serum (FBS, VivaCell, Shanghai, China, C04001) and 1% penicillin/streptomycin (pen/strep). FreeStyle 293F cells (Thermo Fisher Scientific) were cultured in SMM 293-TII Expression Medium (Sino Biological Inc., M293TII). All cells were maintained in a 37°C incubator at 5% CO2

### Protein expression and purification

The genes encoding the ectodomain of SARS-CoV-2 spike including WT and XBB.1.5 were constructed with foldon trimerization motif, His tag, tandem strep-tag II and FLAG tag at C-terminal. The genes encoding the S1, NTD and RBD of WT and XBB.1.5 were constructed with a strep-tag II at C-terminal. The genes encoding the S2 of WT and XBB.1.5 were constructed with a His tag at C-terminal. Spike trimer, S1, S2, NTD and RBD were expressed in the FreeStyle 293F cells and purified by Streptactin Agarose Resin 4FF (Yeasen, 20495ES60) or Ni-NTA Agarose (Qiagen, 30210).

### Spike-specific Single B cell sorting and PCR

PBMCs were incubated with 200nM XBB.1.5 spike for 30 min at 4°C. After wash, they were stained with cell-surface antibodies: CD3-BV510 (BioLegend, 317331), CD19-PE/Cy7 (BioLegend, 302215), CD27-APC (BioLegend, 356409), CD38-APC/Cy7 (BioLegend, 356615), human IgM-AF700 (BioLegend, 314537), human IgD-perCP/Cy5.5 (BioLegend, 348207), anti-His-AF488 (Proteintech, CL488-66005), anti-FLAG-PE (BioLegend, 637309) and DAPI. The stained cells were washed with FACS buffer (PBS containing 2% FBS) and resuspended in 500μl FACS buffer. Spike-specific single B cells were gated as DAPI^-^CD3^−^CD19^+^CD27^+^IgD^-^His^+^FLAG^+^ and sorted into 96-well PCR plates containing 4μl lysis buffer (0.5×PBS, 0.1M DTT and RNase inhibitor) per well. After reverse transcription reaction, variable regions of heavy and light chains were amplified by nested PCR and cloned into human IgG1 expression vectors. Plasmids for paired heavy and light chains were co-transfected into HEK293T cells or FreeStyle 293F cells. Antibodies were purified with Protein A magnetic beads (GenScript, L00273).

### ELISA

Spike, S1, S2, NTD or RBD were coated onto 96-well ELISA plates (100 ng/well) and incubated at 4°C overnight. After blocking with PBS containing 10% FBS, HEK293T supernatant was added to the wells and incubated at 37°C for 1hr. HRP-conjugated goat anti-human IgG antibodies (Zen-bio, 550004; 1:2000 dilution) were added to the wells and incubated at 37°C for 1hr. TMB substrate (Sangon Biotech, E661007-0100) was added to the wells and incubated at room temperature for 5 mins. The reaction was stopped by TMB Stop Solution (Sangon Biotech, E661006-0500) and absorbance at 450 nm was measured.

### Pseudovirus neutralization assay

To generate pseudoviruses, HEK293T cells were transfected with psPAX2, pLenti-luciferase and spike-encoding plasmids using polyetherimide (PEI). Supernatants with pseudoviruses were collected 48hrs after transfection. 3-fold serially diluted plasma (starting at 1:20), HEK293T supernatant (starting at 1:4.5), or mAbs (starting at 10μg/ml) were mixed with pseudoviruses at 37°C for 1hr. HEK293T-hACE2 cells (1.5×10^4^ per well) were added into the mixture and incubated at 37°C for 48hrs. Cells were lysed to measure luciferase activity by Bright-Lite Luciferase Assay System (Vazyme Biotech, DD1204-02). The percentages of neutralization were determined by comparing with the virus control. The plasmids encoding spike of WT, Alpha, Beta, Gamma, Delta, BA.1, BA.4/5, XBB, XBB.1.5 and SARS-CoV-1 were kindly provided by Prof. Linqi Zhang at Tsinghua University. The plasmids encoding spike of BA.2.75 was kindly provided by Prof. Zezhong Liu at Fudan University. The sequences of EG.5, EG.5.1, FL.1.5, FL.1.5.1, HK.3, BA.2.86 and JN.1 spike were synthesized and cloned into expression vectors.

### Competition ELISA

NTD-specific and RBD-specific neutralizing mAbs were labeled with HRP (ProteinTech, PK20001). XBB.1.5 spike was coated onto 96-well ELISA plates (50 ng/well) and incubated at 4°C overnight. After blocking with PBS containing 10% FBS, blocking mAbs were added to the wells and incubated at 37°C for 1hr. HRP-labeled detecting mAbs were added to the wells in the presence of competing mAbs. The ratio of blocking antibody and detecting antibody is 100:1. After 1hr incubation at 37°C, TMB substrate (Sangon Biotech, E661007-0100) was added to the wells and incubated at room temperature for 30 mins. The reaction was stopped by TMB Stop Solution (Sangon Biotech, E661006-0500) and absorbance at 450 nm were measured. The percentages of signal decrease caused by blocking mAbs were calculated.

### ACE2 competition FACS

5nM biotinylated XBB.1.5 spike was incubated with 10μg/ml competing mAbs for 30min at 4°C, then the mixtures were added to HEK293T-hACE2 cells. After 30min at 4°C, the cells were washed with FACS buffer. Streptavidin-APC (BioLegend, 405207) and DAPI were added to the wells at 4°C for 30min. The stained cells were washed with FACS buffer and resuspended in 200μl FACS buffer. Positive cells were gated as DAPI^-^XBB.1.5 spike^+^. The percentages of signal decrease caused by competing mAbs were calculated.

### Germline antibodies binding FACS

The plasmids encoding spikes of SARS-CoV-2 WT, Alpha, Beta, Gamma, Delta, BA.1, BA.4/5, BF.7, BA.2.75, BQ.1, XBB.1.5, EG.5.1, BA.2.86 were transfected into HEK293T cells. After 48hr, cells were harvested and incubated with 50μg/ml germline antibodies for 30min at 4°C. After washed with FACS buffer, cells were stained with goat anti-human IgG FITC (Proteintech, SA00003-12) and DAPI. The stained cells were washed with FACS buffer and resuspended in 200μl FACS buffer. Positive cells were gated as DAPI^-^IgG^+^.

## Acknowledgements

The authors thank all donors for providing peripheral blood. This work was supported by the National Key Plan for Scientific Research and Development of China (Grant No. 2022YFC2305800) and the National Natural Science Foundation of China (Grant No. 32200765, Grant No. 32370941).

---

**Figure S1.**
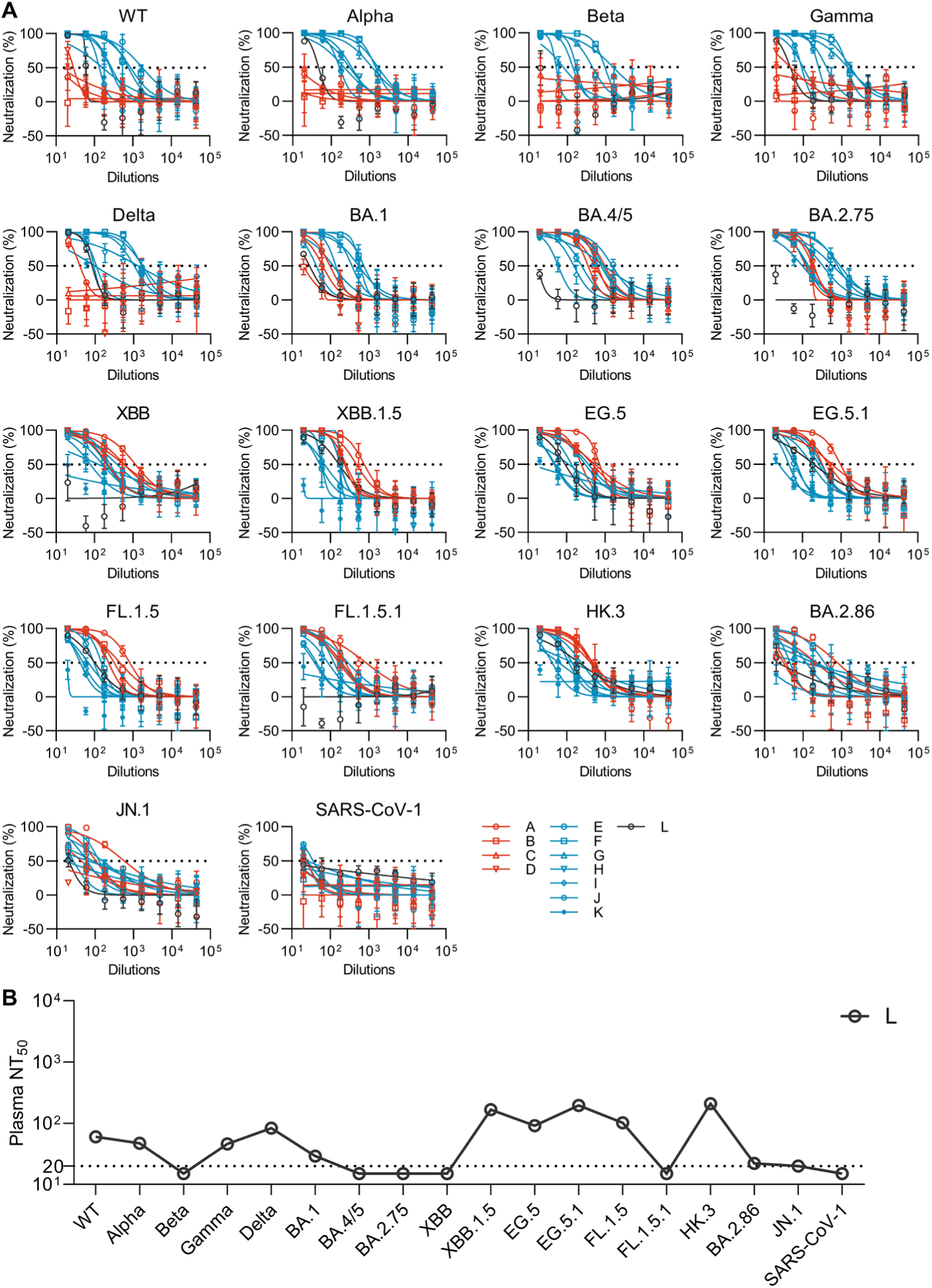
Plasma neutralizing antibody titers. (A) Measurement of neutralizing antibody titers against a panel of pseudoviruses. Data are represented as mean ± SD. (B) Neutralizing antibody titers of donor L.

**Figure S2.**
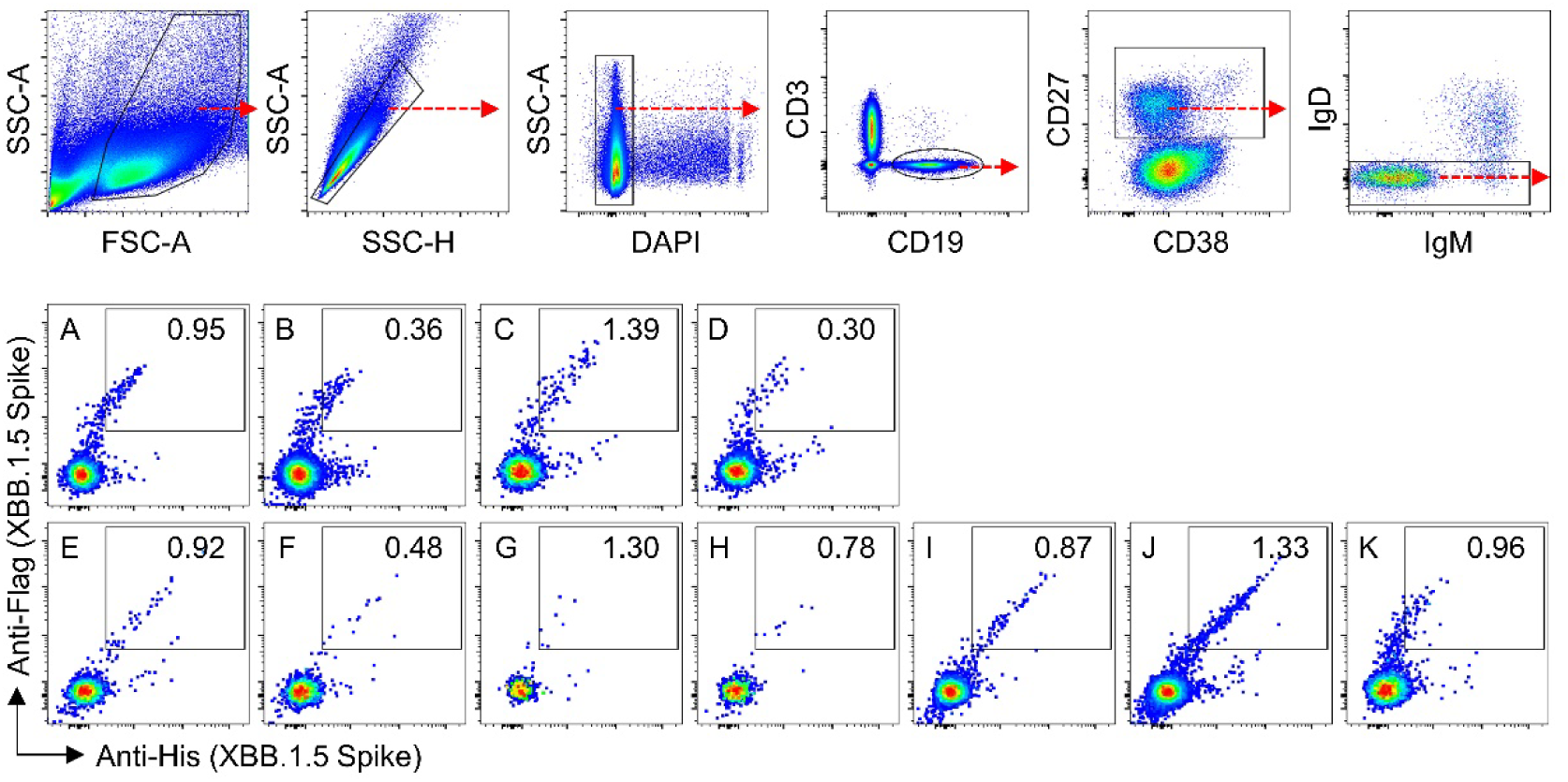
FACS plots representing gating strategies for XBB.1.5 spike-specific single B cell sorting.

**Figure S3.**
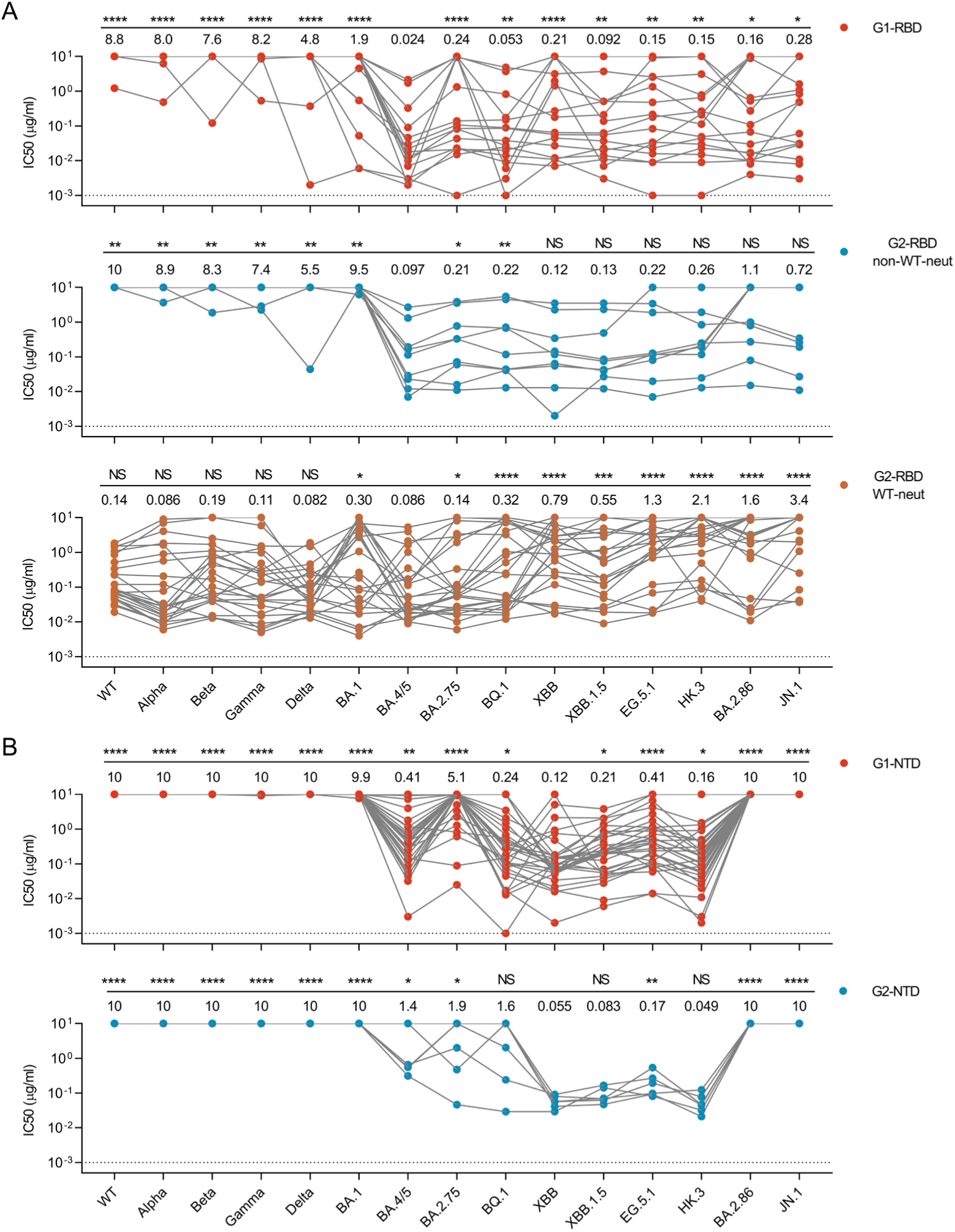
Neutralizing potency of mAbs against different variants. (A) IC50s of RBD-specific mAbs. The numbers below the line are geometric mean IC50 of mAbs against the pseudoviruses. IC50s against BA.4/5 are compared with other variants. (B) IC50s of NTD-specific mAbs. The numbers below the line are geometric mean IC50 of mAbs against the pseudoviruses. IC50s against XBB are compared other variants. Statistical analysis was performed by two-tailed Wilcoxon matched-pairs signed rank test. NS (Not significant) p > 0.05, *P < 0.05, **P < 0.01, ***P < 0.001, ****P < 0.0001.

**Table S1.**
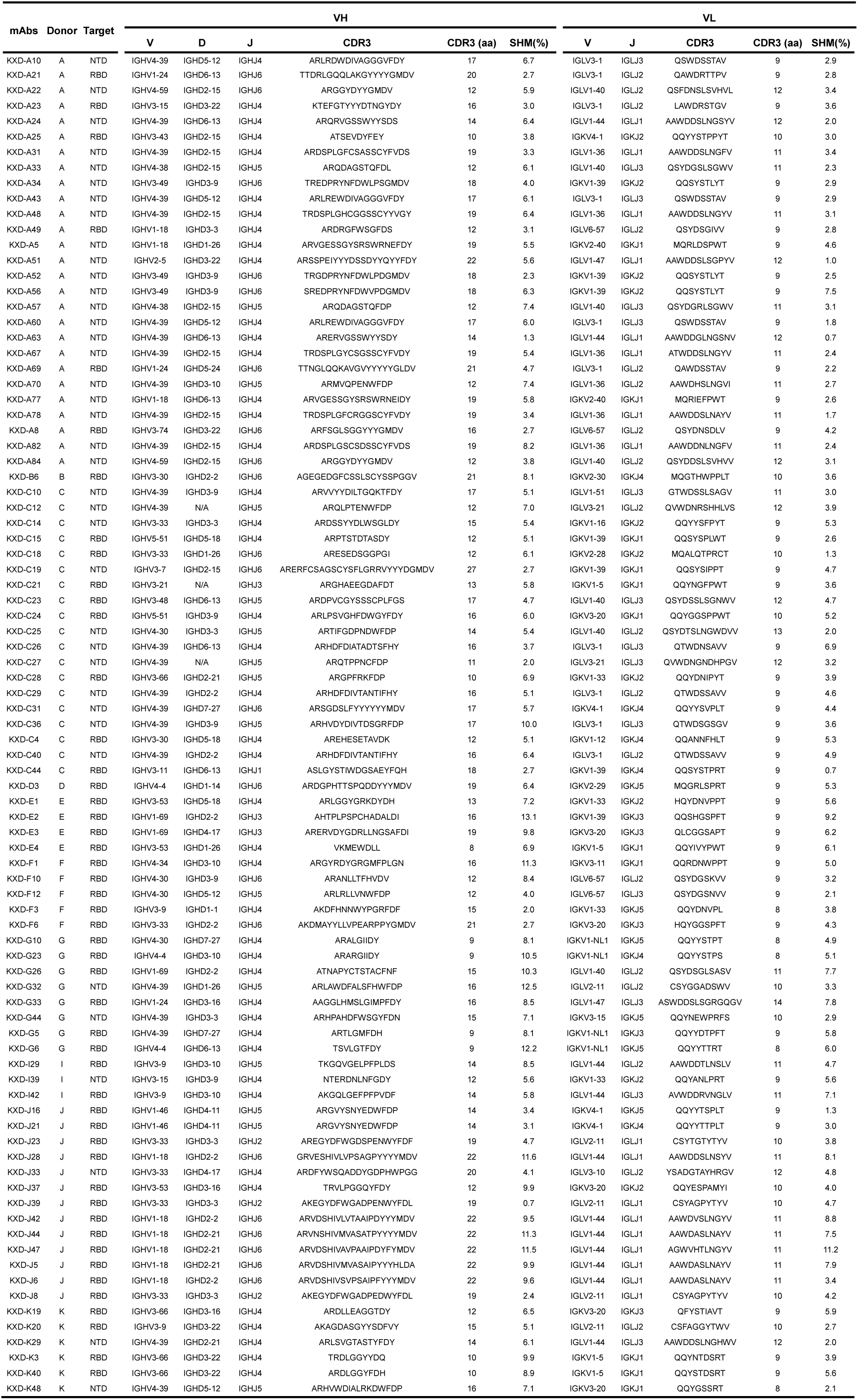
Origins, targets and sequence information of 87 neutralizing mAbs.

